# Repeat *Ascaris* challenge reduces worm intensity through gastric cellular reprograming

**DOI:** 10.1101/2024.08.29.610358

**Authors:** Yifan Wu, Charlie Suarez-Reyes, Alexander R. Kneubehl, Jill E. Weatherhead

## Abstract

Ascariasis (roundworm) is the most prevalent parasitic nematode infection worldwide, impacting approximately 500 million people predominantly in low- and middle-income countries (LMICs). While people of all ages are infected with *Ascaris*, infection intensity (defined by worm burden) paradoxically peaks in pre-school and school aged children but then declines with age. The cause of age-dependent *Ascaris* worm intensity is not well understood but may be dependent on cellular changes in mucosal barrier sites. We have previously found that the gastric mucosa is a critical barrier site for *Ascaris* infection. Following oral ingestion of *Ascaris* eggs, larvae use AMCase secreted by gastric chief cells and acid secreted by gastric parietal cells to hatch. Once hatched, larvae translocate across the gastric mucosa to initiate the larval migratory cycle. However, inducing mucosal injury with administration of Tamoxifen induces mucosa cellular changes that impairs *Ascaris* hatching and reduces larval translocation across the gastric mucosa. In this study we established a repeated *Ascaris suum* challenge mouse model and evaluated if repeated Ascaris challenge also lead to cellular changes in the gastric mucosal barrier. We found that repeated *Ascaris* challenge caused cellular changes in the gastric mucosa which reduced worm intensity in the liver independent of the adaptive immune response. Thus, in endemic regions, where individuals experience recurrent infection throughout their lives, gastric cellular changes may be a key mechanism leading to the observed age-dependent *Ascaris* worm intensity changes from childhood to adulthood.

## Introduction

Ascariasis (roundworm) is the most prevalent parasitic nematode infection worldwide, impacting approximately 500 million people (1, 2). In endemic regions, particularly in low and middle income countries (LMICs), people are infected with either *Ascaris lumbricoides* or *Ascaris suum* by oral ingestion of eggs from contaminated soil or water (3-6). After ingestion, *Ascaris* larvae hatch in the gastrointestinal tract and initiate a transient, highly immunogenic, larval migration cycle through the host’s liver, lungs, and eventually the intestines to develop into adult worms where they live for up to 2 years (7). In endemic regions, individuals experience recurrent infection throughout their lives (8). While people of all ages are are infected with *Ascaris*, infection intensity (defined by worm burden) paradoxically peaks in pre-school and school aged children but then declines with age.

The cause of age-dependent *Ascaris* worm intensity is not well understood but likely reflects changes in host behavior as children age into adulthood as well as changes to the adaptive immune response and/or cellular changes at mucosa barrier sites following repetitive infection. Specifically, because *Ascaris* is transmitted by oral ingestion of *Ascaris* eggs found in the environment, childhood behaviors such as pica (also known as geophagia) put younger children at risk of high worm intensity (9). Additionally, adaptive immunity such as activation of CD4^+^ T cells (10) as well as IgE isotype class switching (11) may provide protective immunity against *Ascaris* at different immune barrier sites following recurrent infection. In a mouse model of *Ascaris* larval migration, recurrent infection was associated with reduced worm intensity in the lungs and was associated with increased number of T cells, in addition to innate immune cells such as neutrophils, eosinophils and macrophages (10). While each of these mechanisms are likely contributing to age-dependent *Ascaris* worm intensity, none fully explain observed age-dependent intensity changes. Thus, it is important to evaluate cellular changes at mucosa barrier sites which may better explain the decrease in worm intensity as children age to adulthood but have largely been unexplored.

The gastric mucosa is a critical barrier site for *Ascaris* infection. We have previously shown that following oral ingestion of *Ascaris* eggs, larvae use AMCase secreted by gastric chief cells and acid secreted by gastric parietal cells to hatch. Despite prior literature suggesting larvae hatch in the intestines (12), we recently showed evidence for hatching in the host stomach (13). Once hatched, larvae translocate across the gastric mucosa to initiate the larval migratory cycle. However, inducing gastric mucosa cellular changes with administration of Tamoxifen (14) to reduce AMCase and gastric acid impairs *Ascaris* hatching and reduces larval translocation across the gastric mucosa. Further, impairing *Ascaris* translocation across the gastric mucosa reduces worm intensity in the liver and the lungs during the larval migration cycle suggesting a critical role for gastric mucosa in determining worm intensity (13). The aim of this study was therefore to assess the mechanism by which gastric mucosa might contribute to age dependent changes in worm intensity. Towards this goal, we established a repeated *Ascaris suum* challenge mouse model and evaluated the impact on the gastric mucosal barrier. We found that repeated *Ascaris* challenge caused cellular changes in the gastric mucosa which reduced worm intensity in the liver independent of the adaptive immune response.

## Results

### *Repeat Ascaris challenge reduces* Ascaris *worm intensity in the liver*

Prior literature has shown that *Ascaris* worm intensity in mice is reduced in the lungs following repeat *Ascaris* challenge (10). However, the liver is the first sight in the migration cycle. Therefore, we first asked if repeat *Ascaris* challenge in mice impacts larval intensity in the liver. Towards this, we used a repeat *Ascaris* challenge model in which wild-type mice were infected with 2500 embryonated *Ascaris* eggs twice a week for two weeks (a total of 4 challenges) by oral gavage (**Figure 1A**) compared to a single infection with *Ascaris* eggs (15). Four days following the last *Ascaris* egg challenge, mice were euthanized. We found that mice exposed to repeat *Ascaris* challenge had reduced *Ascaris* larval burden in the liver compared to mice exposed to a single *Ascaris* challenge (**Figure 1B**). Additionally, our prior work demonstrated that impairing gastric AMCase and acid secretion inhibited *Ascaris* hatching, larval translocation across the gastric mucosa and migration to the liver and lungs (13). Based on these previous data, the stomach may be a primary mucosa barrier site to prevent *Ascaris* worm intensity. To determine the impact of repeat *Ascaris* challenge on gastric acid and AMCase secretion from the gastric mucosa, we next measured *atp4a*, a gene that encodes the proton pump that pumps out gastric acid, and *chia1*, a gene that encodes AMCase, mRNA expression from stomach tissue harvested from our wild-type repeat *Ascaris* challenge mouse model compared to wild-type, naive mice 4 days following the last *Ascaris* egg challenge. Indeed, *atp4a* and *chia1* mRNA expression levels measured by qPCR were both reduced (**Figure 1C**). Decreased Atp4a and AMCase protein levels were confirmed by western blot (**Figure 1D**). Together these results demonstrate a reduction in *Ascaris* worm intensity in the liver following repeat *Ascaris* challenge that is associated with decreased gastric mucosa function, specifically reduction in gastric acid and AMCase secretion.

**Figure 1:**
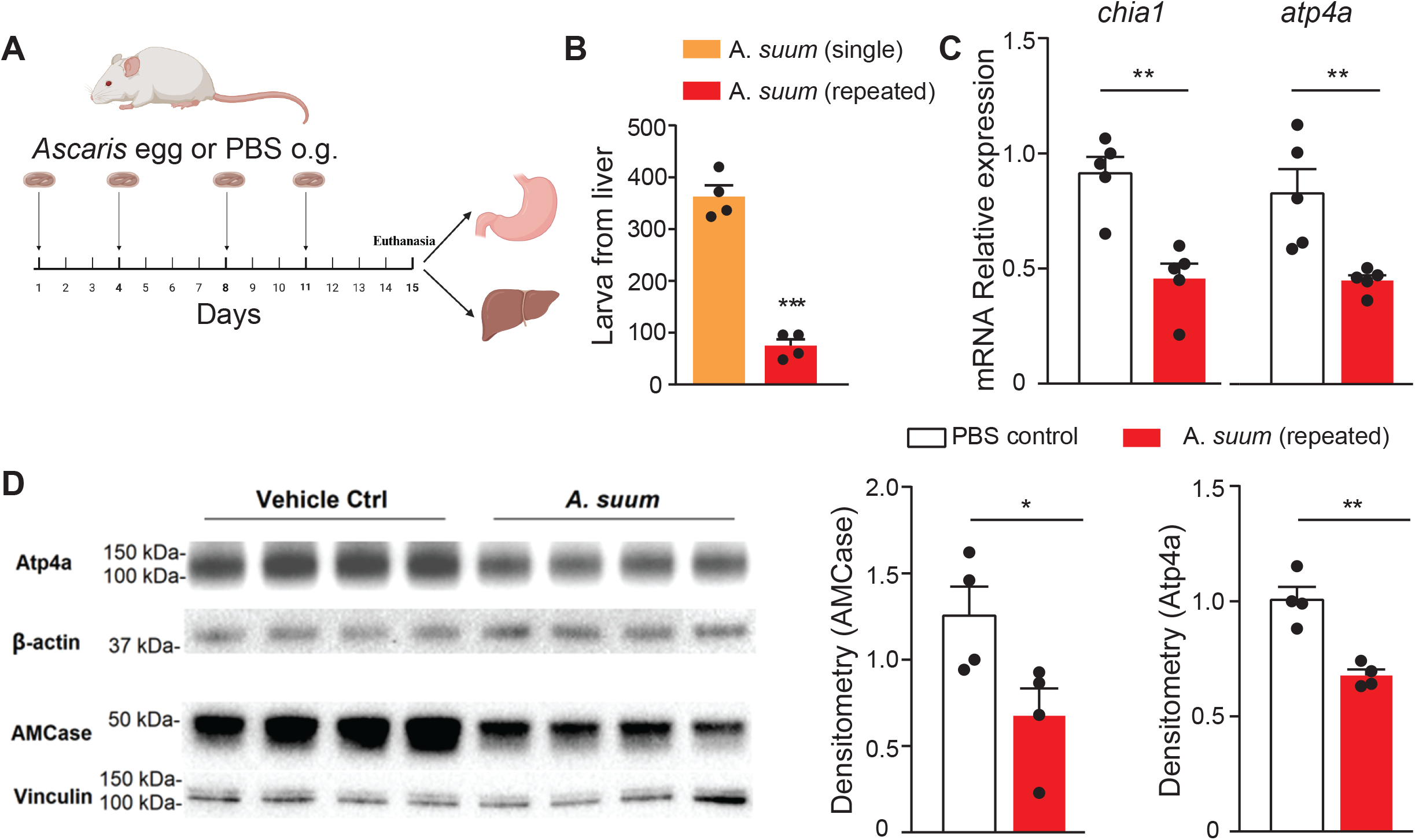
Repeat *Ascaris* challenge by oral gavage reduces *Ascaris* worm intensity in the liver with associated impaired host gastric microenvironment function. Repeat *Ascaris* infection mouse model(A)in which wild-type mice were infected with embryonated *Ascaris* eggs or PBS twice a week for two weeks by oral gavage. Four days following the last infection mice were euthanized and stomach and liver tissue were harvested. Larval count in the liver (B) illustrating decreased intensity for repeat *Ascaris* infection mouse model compared to a single *Ascaris* infection model. mRNA gene expression (C) of *atp4a* (left) and *chia1 (right)* in the stomach by qPCR following repeated *Ascaris* infection compared to PBS, non-infected controls. Western blot with quantification (D) of protein expression of Atp4a and AMCase in the stomach following repeated *Ascaris* infection compared to PBS, non-infected controls.

### *Gastric CD4^+^ T cells are decreased follow repeat* Ascaris *challenge*

Building off our findings of reduced mRNA expression of key gastric genes *atp4* and *chia1*, we next aimed to evaluate changes in the gastric transcriptome following repeat *Ascaris* challenge. Using our wild-type, repeat *Ascaris* challenge model (**Figure 1A**) we compared bulk RNA sequencing from stomach tissue harvested 4 days post last challenge with stomach tissue from naïve mice. A total of 19,166 genes were detected. Considering all genes, stomach tissue harvested from repeat *Ascaris* challenge mice had a distinct transcriptome by euclidean distance matrix (**Supplemental Figure 1A**) and principal component analysis (**Supplemental Figure 1B**). Of those genes, 751 were significantly downregulated and 695 were significantly upregulated in the stomach of wild-type, repeat *Ascaris* challenge wild-type mice compared to wild-type, naive mice as observed by volcano plot (**Supplemental Figure 1C**).

Based on the reported role of the adaptive immune response in controlling *Ascaris* infection (10), we hypothesized that recruitment and activation of CD4^+^ T cells would provide anti-*Ascaris* immunity in the stomach directly by killing the parasite or indirectly by altering gastric acid and AMCase secretion. To evaluate if gastric CD4^+^ T cells provide anti-*Ascaris* immunity following repeat *Ascaris* challenge, we identified T cell recruitment and activation genes in the stomach tissue from our RNA sequencing data. Surprisingly, genes involved in T cell recruitment and activation were downregulated in the stomach following repeat *Ascaris* challenge compared to naïve controls (**Figure 2A**). Common T cell chemokine genes including cxcl13, ccl21b, ccl5, cxcr3, ccr7, il16, xcl1were selected from the transcriptomic analysis for confirmation of mRNA expression by qPCR (**Figure 2B**). Given this result, we next asked if the lack of T cell recruitment and activation chemokine gene expression was secondary to reduced CD4^+^ T cells in stomach tissue. Using flow cytometry on stomach tissue from wild-type, repeat *Ascaris* challenge mice compared to wild-type, naïve controls, we found that repeat *Ascaris* challenge was associated with reduced CD4^+^ T cells in stomach tissue (**Figure 2C**). These data suggest that gastric adaptive immune responses, specifically CD4^+^ T cells, do not play a role in anti-*Ascaris* immunity as the gastric mucosa. To confirm that *Ascaris* larval hatching and translocation is not modulated by the adaptive immune response, particularly CD4^+^ T cells, we challenged mice deficient in T cells and B cells (RAG^-/-^ mice) with 2500 embryonated *Ascaris* eggs twice a week for two weeks (a total of 4 challenges) by oral gavage (**Figure 2D**). Despite deficiency in T cells and B cells, RAG^-/-^, repeat *Ascaris* challenge mice had reduced larvae intensity in the liver compared to RAG^-/-^ mice exposed to a single *Ascaris* challenge (**Figure 2E**). Larva burden in RAG^-/-^, repeat *Ascaris* challenge mice were similar to wild-type, repeat *Ascaris* challenge mice (**Figure 1B**) confirming gastric adaptive immune responses do not play an essential role in reducing larval worm intensity.

**Figure 2:**
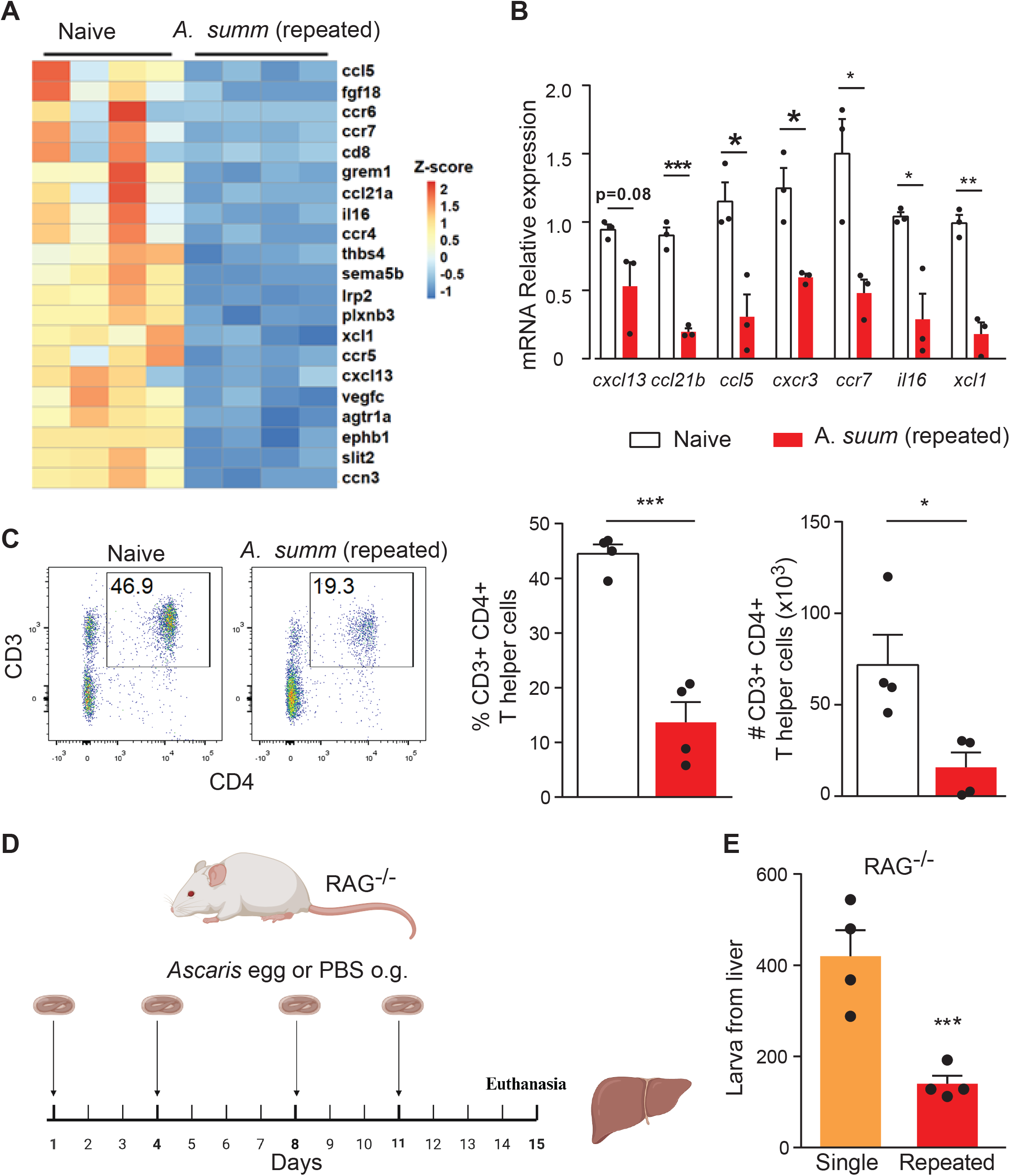
Reduced worm intensity is independent of the gastric adaptive immune response following repeated *Ascaris* challenge. Repeat *Ascaris* infection mouse model in which wild-type mice were infected with embryonated *Ascaris* eggs or PBS twice a week for two weeks by oral gavage. Four days following the last infection mice were euthanized and stomach tissue was harvested. Expression levels of T cell recruitment and activation genes in gastric tissue (A) from bulk RNA sequencing. mRNA concentrations of T cell recruitment and activation genes from A in gastric tissue by qPCR. Flow cytometry with quantification (C) showing concentration of CD4^+^ T cells in gastric tissue. Repeat *Ascaris* infection mouse model (D) in which mice deficient in T and B cells (RAG^-/-^) were infected with embryonated *Ascaris* eggs or PBS twice a week for two weeks by oral gavage. Four days following the last infection mice were euthanized and liver tissue was harvested. Larval count in the liver (E) illustrating decreased intensity for repeat *Ascaris* infection mouse model compared to a single *Ascaris* infection.

### *Repeat* Ascaris *challenge causes gastric cellular reprogramming*

Based on evidence that the gastric adaptive immune response is not responsible for *Ascaris* larvae intensity (**Figure 2E**) and given decreased expression of *atp4a* and *chia1* mRNA expression by qPCR (**Figure 1C**), we hypothesized that the decreased intensity was due to cellular changes in the gastric mucosal barrier. We therefore evaluated our stomach transcriptome data to evaluate genes associated with gastric cellular reprogramming and found decreased expression of parietal cell proton pump genes, atp4a and atp4b (**Figure 3A**). Decreased parietal cell gene expression was accompanied by increased expression of genes involved in apoptosis by bulk RNA sequencing (**Figure 3B**). Given RNA sequencing evidence of increased expression of apoptotic pathways and decreased parietal cell function, we evaluated gastric histopathology from wild-type, repeat *Ascaris* challenge mice compared to wild-type, naïve mice (**Figure 3C**). We found repeat *Ascaris* challenged mice had increased apoptotic bodies in the gastric tissue consistent with parietal cell loss (**Figure 3D**).

**Figure 3:**
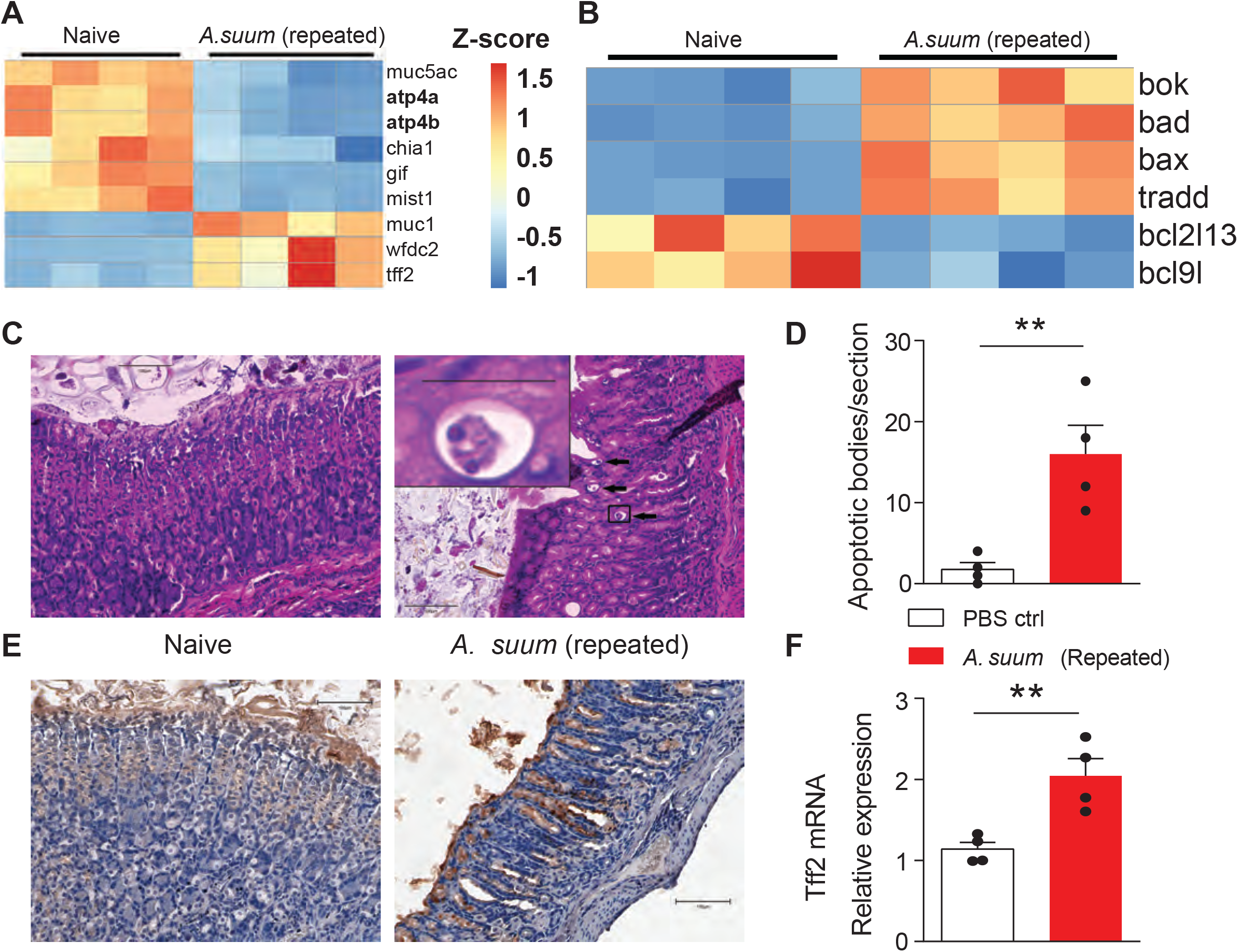
Repeated *Ascaris* challenge causes cellular changes to the gastric mucosa. Bulk RNA sequencing gene expression changes (A) for genes in stomach tissue associated with cellular reprogramming following mucosal injury showing decreased expression of atp4a and atp4b and increased expression of ttf2. Bulk RNA sequencing gene expression changes (B) in stomach tissue also reveal increased expression of genes associated with cellular apoptosis. Immunohistochemistry (C) showing increased apoptotic bodies (insert) consistent with parietal cell death identified by histopathology (H&E staining) in wild-type repeat *Ascaris* challenge model compared to wild-type naive mice. Quantification of apoptotic bodies (F) in the gastric mucosa showing increased cell death in gastric mucosa following repeat *Ascaris* challenge compared to naive mice. Immunohistochemistry (E) showing increased expression of ttf2 (brown staining) and qPCR (F) quantification of increased ttf2 mRNA expression in gastric mucosa in repeated *Ascaris* challenge model compared to naive mice.

Based on decreased *chia1* mRNA expression we hypothesized that chief cells were undergoing transdifferentiation, Indeed, stomach transcriptomic data demonstrated increased gene expression of trefoil factor 2 (*ttf2*) also known as spasmolytic polypeptide and *muc1* as well as decreased gene expression of *mist1* (**Figure 3A**) following repeat *Ascaris* challenge. Increased expression of *ttf2* suggests gastric chief cells are transdifferentiating into spasmolytic polypeptide-expressing metaplasia (SPEM) which classically occurs following parietal cell loss. Upregulation of *ttf2* was visualized in the gastric crypts by immunohistochemistry staining (**Figure 3E**) and quantitated by RNA expression using qPCR (**Figure 3F)**. Together these data demonstrate repeat *Ascaris* challenge induces cellular reprogramming of the gastric mucosa including apoptosis of parietal cells and transdifferentiation of chief cells suggesting a novel mechanism for age dependent *Ascaris* intensity differences.

## Discussion

We found that repeat *Ascaris* infection reduces worm intensity through gastric cell reprograming, specifically parietal cell apoptosis and chief cell transdifferentiation into SPEM cells. These gastric cellular changes resulted in reduced gastric acid and AMCase secretion in the stomach. The lack of gastric acid and AMCase in the stomach impaired *Ascaris* egg hatching and translocation across the gastric mucosa, resulting in reduce larval intensity in the liver. Interestingly, these changes occurred independent of the adaptive immune response. Together, these findings support the role of the gastric mucosa as a primary barrier site in the prevention of *Ascaris* following repeat infection.

Our finding that repeat challenge with *Ascaris* induces the host gastric mucosa to undergo cellular changes which reduce AMCase and, thus, prevent *Ascaris* infection, raises the possibility that gastric cellular changes may be a conserved anti-nematode mechanism to reduce parasite burden in the host. Nematode eggs are covered in a thick chitin exterior that protects the egg in harsh environments but conversely require a mechanism within the host to break down this chitin exterior in order to facilitate infection. We have previously shown for *Ascaris* infection that gastric AMCase serves this role, breaking down the chitinous egg (13). Due to similar egg composition amongst different nematodes species, it is reasonable to hypothesize that gastric cellular changes are a conserved mechanism in the host to reduce nematode intensity. Future studies can assess the role of gastric AMCase on worm intensity for other nematodes. If true, this mechanism could be targeted for the development of a pan-nematode vaccine which has been hampered by the complexity of the nematode life cycle and the host systemic immune response to the nematode. Identifying a gastric mucosal target for vaccine development could overcome the limitations of a vaccine that causes a system type-2 immunity response which have hampered nematode vaccine development to date.

Gastric mucosal injury triggers the development of pyloric metaplasia to create an environment that rapidly protects and restore the mucosal barrier. Interestingly, these cellular changes also lead to overexpression of surface receptors, such as selectins like sialyl-Lewis^X^ receptor and integrins, used by *Helicobacter pylori* to adhere to and colonize the gastric mucosa (16, 17). Adherence to the gastric corpus epithelium allows *H. pylori* colonization and expansion of pyloric metaplasia in the gastric corpus in a positive feedback loop^8^. The type of gastric cellular changes we observed following *Ascaris* infection would therefore suggest that individuals who have endured repeat *Ascaris* infection and subsequent *Ascaris*-induced pyloric metaplasia would have a higher risk for *H. pylori* colonization. Indeed, the geographic niche of these pathogens overlaps. However, no studies to date have examined co-infection. Interestingly, compared to children living in high-income countries, those living in LMICs are more commonly infected with *H*.*pylori* in early childhood (18, 19). By adulthood, up to 50-80% of individuals in LMICs are colonized with *H*.*pylori* compared to <40% of individuals in high-income countries (20, 21). The reasoning behind the earlier colonization and more prevalent disease in LMICs remains unknown. However, given the significant geographic overlap between *H. pylori* and *Ascaris*, co-infection with these pathogens is likely and development of *Ascaris*-induced pyloric metaplasia at an early age may influence the lifetime risk of *H. pylori*.

Based on our findings, it is likely that pyloric metaplasia not only aids in gastric repair and restoration following mucosal injury but also serves as an Anti-*Ascaris* mechanism to prevent high worm intensity following repeated *Ascaris* infection over time. Thus, in endemic regions, where individuals experience recurrent infection throughout their lives, pyloric metaplasia is a likely contributor to the observed age-dependent *Ascaris* worm intensity changes from childhood to adulthood. However, while *Ascaris*-induced pyloric metaplasia prevents high larval intensity following repeat *Ascaris* infection, pyloric metaplasia may also lead to collateral gastric pathology. Further evaluation of the collateral damage to the gastric mucosa and the down-stream sequalae to the gastrointestinal tract as a result of *Ascaris*-induced pyloric metaplasia is critical to understanding the complex relationship between *Ascaris* and the gastric mucosa barrier.

## Methods

### Mice

Ten 8-week-old BALB/c female mice (wildtype or Rag2^-/-^) were purchased from Jackson Laboratories (cat: 000651) or Taconic Biosciences (cat: 601). Upon arrival, complete randomization of mice into longitudinal groups was performed. Only female mice were used to ensure consistency in infectious burden (22). All mice were housed in a vivarium under specific-pathogen-free conditions. All experimental protocols were approved by the Institutional Animal Care and Use Committee of Baylor College of Medicine and followed federal guidelines.

### *A. suum* experimental murine model

*A. suum* eggs were obtained from adult female worms from infected pigs in the Weatherhead laboratory at Baylor College of Medicine. Briefly, adult female worms were isolated and dissected to remove the uterus. The uterus was then strained through a filter to release unembryonated eggs. The eggs were washed with PBS three times and subsequently resuspended in sulfuric acid for 60 days to allow for embryonation. For single infections, BALB/c mice were treated with a single inoculum of 2,500 embryonated *A. suum* eggs via oral gavage or PBS as previously described (23, 24). For repeated infections, BALB/c mice were infected with an inoculum of 2,500 embryonated *A. suum* eggs or PBS via oral gavage twice per week for 2 weeks. The infectious dose of *Ascaris* has been standardized in the literature in order to replicate human disease in a murine model (23, 25). The *A. suum* life cycle in a murine model mimics the life cycle in humans and has been previously described (23, 26). Following oral gavage of *A. suum* eggs or PBS, mice were euthanized at 4 days post last infection (p.i). and stomachs and liver were harvested. in preparation for experiments described below. For stomach tissue, the forestomach was removed and discarded and the remaining tissue was washed in PBS.

### Quantitative PCR

Stomach tissue was homogenized in TRIzol (15596026, Thermofisher scientific, Waltham MA) using M tubes on gentleMACS dissociator (130-093-236, Miltenyi, Bergisch Gladbach, Germany) for RNA extraction. Relative expression of mRNA for *Chia1* and *Atp4a* to 18s was detected by two-step, real-time quantitative reverse transcription-polymerase chain reaction (RT-PCR) with the 7500 Real-Time PCR System (Applied Biosystems, Foster City, CA) using Taqman probe (Mm00458221, Mm00444417, Hs03003631 Invitrogen, Carlsbad, CA)(27, 28).

### Western blot

Stomach tissue was homogenized in tissue lysis buffer (50mM NaCl, 20mM HEPES, 1mM EDTA, 2% Triton-X 100, 10% glycerol, with proteinase and phosphatase inhibitor) for protein extraction (29) Protein expression level of AMCase and Atp4a was detected using western blot. (4%-12% Nupage Bis-Tris gel, Thermo Fisher Scientific, Waltham MA)(Antibody: ab207169, ab174293, 1:2000, Abcam, Cambridge MA; #4970S, #13901T, 1:2000 Cell signaling, Danvers MA).

### Histopathology

Gastric tissue was fixed in 10% neutral-buffered formalin solution, processed and embedded in paraffin. 5 μm sections were cut and slides were stained with H&E. Apoptotic bodies were numerated from each section (We only have 4 sections made). Alternatively, gastric tissue was processed and embedded in paraffin, 5 μm sections were cut and immunohistochemistry was completed. Briefly, slides were deparaffinized, then permeabilized using 0.2% Triton X 100 (X100, Sigma-Aldrich, St. Louis, MO). After that, antigen recovery was completed using Diva Decloaker (DV2004, Biocare Medical, Pacheco, CA), and peroxidase blocked using 3% hydrogen peroxide. Slides were then blocked using 5% bovine serum albumin (A0100-005, Gendepot, Baker, TX) for 1 hour at room temperature, and incubated with primary anti-Tff2 antibody in 5% bovine serum albumin (1:50, 13681-1-AP, Thermofisher scientific, Waltham MA) overnight at 4 °C. Slides were then incubated and stained using ABC-HRP and DAB kit (PK-6200, SK-4100, Vector Laboratories, Newark, CA). Slides were then counterstained with hematoxylin, dehydrated and mounted.

### Isolation of *A. suum* larva from the liver

Mice infected with *A. suum* eggs were euthanized 4 days p.i. to assess larval burden in the liver. We measured larval burden in the liver at day 4 p.i. because this is when intensity peaks (15). Liver tissue was harvested and macerated with scissors, suspended into pre-warmed PBS and transferred in a modified Baermann apparatus. The collection system was then filled up to 40 ml of pre-warmed PBS and incubated at 37°C for 4 hours. Following incubation, the solution containing larvae was collected from the apparatus, centrifuged at 800 xg for 5 minutes at room temperature, and washed with water to remove red blood cells. Larvae were washed with PBS for additional 3 times and counted under the microscope.

### Bulk RNA sequencing

Stomach tissues from mice post multiple infections were collected 4 days p.i. into 1mL of 1x DNA/RNA Shield reagent (R1100-50, Zymo Research, Irvine, CA) in a 2mL ZR bashingbead lysis tube (S6012-50, Zymo Research, Irvine, CA). The tissues were lysed using a Precellys 24 homogenizer (03119.200.RD000, Bertin Technologies, Montigny-le-Bretonneux, France) using the 6m/sec setting for 40s. The homogenates were transferred to 1.5mL DNA LoBind tubes (022431021, Eppendorf, Hamburg, Germany) and shipped on dry ice to SeqCenter (Pittsburgh, PA) for RNA extraction, polyA purification, and RNA sequencing. RNA was extracted using the Quick-RNA miniprep kit (R1054, Zymo Research, Irvine, CA). PolyA selection and library preparation for RNA sequencing were performed using the Illumina stranded mRNA prep kit, 2×150bp reads (20040534, Illumina, San Diego, CA). RNA sequencing was performed on a NovaSeq X Plus instrument. Each sample was sequenced using SeqCenter’s 50M paired end, polyA read option and on average 20 Gbp of read data were generated for each sample. Read data were submitted to the Sequence Read Archive (BioProject PRJNA1141177).

RNA sequencing analysis was performed using Nextflow’s nf-core/rnaseq pipeline (v3.14.0) (1) to generate read counts using the STAR aligner (2) and salmon (3) for gene counts. Differential gene expression analysis was performed using DEseq2 (4). Exploratory data analysis from DEseq2 (PCA and Euclidean distance matrix analysis) indicated two outlier samples, one in each group. The DEseq2 analysis was performed again with these outliers removed and all further analysis was done without the outliers. To visualize the differentially expressed genes, a volcano plot was generated using the EnhancedVolcano R package (5). Gene ontology (GO) analysis was performed using clusterProfiler R package (6). Heatmaps of gene expression for selected genes were generated using the pheatmap R package (7). Analytical scripts along with version information and the R environment used can be found on GitHub (https://github.com/kneubehl/repeated-ascaris-infection-gastric-rnaseq-analysis). The DEseq2 normalized counts for each sample were submitted to the Gene Expression Omnibus repository and are publicly available (PRJNA1141177).

### Single Cell suspension

Stomach tissues from mice post multiple infections were collected 4 days p.i. Stomachs were cut into small pieces and incubated in digestion buffer (2mg/ml collagenase (#LS004177, Worthington), 0.04mg/ml DNAse (#10104159001, Sigma) 1, 20% FBS in HBSS) for 1 h at 37°C after which they were deaggregated by pressing through a 40 μM nylon mesh and centrifuged at 400 × g for 5 minutes at 4°C. Supernatants were discarded, and 1.5 mL of ACK (Thermofisher scientific, Waltham MA) was added and incubated for 3 min at room temperature for erythrocyte lysis. ACK was then neutralized with 7.5 mL of complete RPMI-1640 (Corning, NY), with 10% FBS and 1% Pen Strep, Gibco, Waltham MA). The resulting single cell suspension was prepared for flow cytometry analysis(30).

### Flow cytometry

Total stomach cells isolated above were stained with Live/Dead Fix Blue (L34961, Thermofisher scientific, Waltham MA) and CD45 (103112, Biolegend, San Diego, CA). For T helper cell staining, cells were stained with CD3, CD4 (100222, 100412, Biolegend, San Diego, CA). Cells were then separated into 3 individual groups, permeabilized/fixed using Transcription Factor Buffer Set (562574, BD Biosciences, San Jose, CA), and stained individually for T-bet, GATA3 or RORgt (561265, 560074, 562607, BD Biosciences, San Jose, CA). For macrophages and eosinophils, cells were stained with F4/80 and siglec F. (157309, 155506, Biolegend, San Diego, CA). For ILC2s, Cells were stained with lineage cocktail (133310, Biolegend, San Diego, CA) and then permeabilized/fixed using Transcription Factor Buffer Set (562574, BD Biosciences, San Jose, CA), and stained for GATA3 (560074, BD Biosciences, San Jose, CA) (30, 31).

## Statistical analysis

Data are presented as means ± standard errors of the means. Significant differences relative to PBS-challenged mice are expressed by P values of <0.05, as measured two tailed Student’s t-test, one-way or two-way ANOVA followed by Tukey’s test for multiple comparison. Data normality was confirmed using the Shapiro-Wilk test. Experiments were completed in duplicate.

## Data availability

The data that support the findings of this study are available from the corresponding authors upon reasonable request. Sequencing data and RNAseq analysis is publicly available on NCBI through BioProject PRJNA1141177. Analytical scripts used in the analysis of the sequencing data are publicly available on GitHub at https://github.com/kneubehl/repeated-ascaris-infection-gastric-rnaseq-analysis.

## Acknowledgement

This manuscript was prepared with the assistance of a science writer, Ariel M Lyons-Warren. Funding for the manuscript was provided by NIH NIAID K08A143968 and the Pediatric Infectious Disease Society Foundation Pichichero Family Foundation Vaccines for Children Initiative Research Award. Alexander R. Kneubehl was supported through the Infection and Immunity T32 Fellowship T32AI055413 at Baylor College of Medicine.

---

**Supplemental Figure 1: Repeated *Ascaris* challenge is associated with distinct gastric transcriptomic profile** Heat map (A) and principal component plot (B) compared to naive gastric tissue. Volcano plot (C) with increased and decreased expression of genes in the gastric mucosa of mice repeatedly challenged with *Ascaris* compared to naïve mice.

