## Supplementary figures and images for "Repeat *Ascaris* challenge reduces worm intensity through gastric cellular reprograming"

### Supplemental Figure 1

**Supplemental Figure 1**

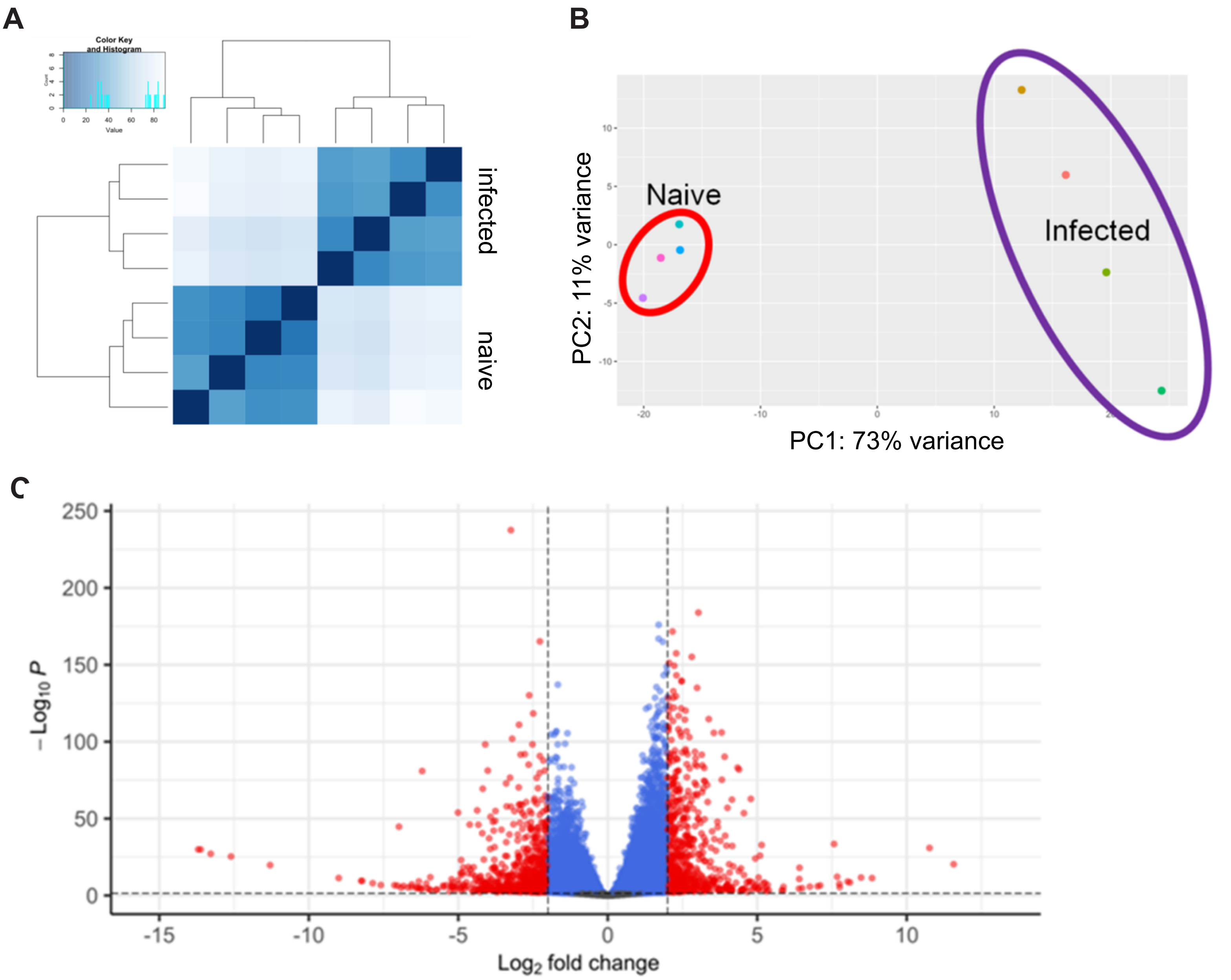
